# Phylogenomics supports monophyly of marsupial crustaceans: a journey to direct development

**DOI:** 10.1101/2025.11.06.686749

**Authors:** Anna-Chiara Barta, Markus Grams, Heather Bracken-Grissom, Saskia Brix, Lívia M. Cordeiro, Brittany Cummings, Stormie Collins, William J. Farris, Sarah Gerken, Christoph G. Höpel, Anne-Nina Lörz, Siena McKim, Kenneth Meland, Luise Kruckenhauser, Jørgen Olesen, Pedro A. Peres, Stefan Richter, Regina Wetzer, Jason Williams, Kevin M. Kocot, Martin Schwentner

## Abstract

Peracarida (marsupial crustaceans) represent one of the most diverse and ecologically important crustacean groups, yet their evolutionary relationships have long been debated. Here, we present the most comprehensive phylogenomic analysis of Peracarida to date, incorporating the relict taxa Thermosbaenacea, Mictacea, Ingolfiellida, and Spelaeogriphacea for the first time in a phylogenomic framework. Our results robustly confirm peracarid monophyly and recover a well-supported clade uniting Mancoida (Isopoda, Tanaidaca, Cumacea), Mictacea, and Spelaeogriphacea. We propose the new taxon Panmancoida to encompass this expanded lineage defined by shared developmental and morphological traits. The inferred phylogeny further suggests that peracarid evolution involved a transition from an intermediate “pseudodirect” developmental mode to the direct development seen in most lineages. We further show that the shift to extensive brood care within the marsupium, accompanied by the loss of a free-swimming larval stage, may have accelerated rates of molecular evolution across lineages. Together, these findings provide a robust evolutionary framework for this major malacostracan lineage and highlight how key reproductive innovations reshaped the genomic and life-history trajectories of the marsupial crustaceans.

## Introduction

Peracarida (marsupial crustaceans) is one of the two major lineages of malacostracan crustaceans, rivaling Decapoda (shrimps, crabs, lobsters, and relatives) in both diversity and ecological scope. With ∼26,400 described species (WoRMS, accessed August 2025), they occupy nearly every marine and many freshwater and terrestrial habitats, where they are often abundant and have an important role in nutrient cycling. The group includes species-rich orders such as Amphipoda, Isopoda, Cumacea, Tanaidacea, and Mysida, as well as several relict lineages with only a handful of species. These latter groups—Bochusacea (6 species), Ingolfiellida (54 species), Lophogastrida (53 species), Mictacea (1 species), Spelaeogriphacea (4 species), Stygiomysida (16 species), and Thermosbaenacea (45 species)—are largely confined to extreme or isolated environments such as caves, subterranean aquifers, and the deep sea. Despite their ecological importance and diversity, phylogenetic relationships among the peracarid orders remain poorly understood, and even peracarid monophyly has been doubted (e.g., Meland & Willasen 2007), leaving fundamental questions about the group’s evolutionary history unanswered.

Traditional morphology-based phylogenetic investigations generally support peracarid monophyly, based on shared derived characters, such as the brood pouch (marsupium; Richter & Scholtz 2001; Poore 2005; Wills et al. 2009; Wirkner & Richter 2010; Grams et al. 2025). However, molecular studies have yielded conflicting results regarding peracarid monophyly. Single-gene analyses based on 18S or 28S rDNA (e.g., Jarman et al. 2000; Spears et al. 2005; Meland & Willassen 2007; Jenner et al. 2009) consistently excluded Mysida from Peracarida. These studies recovered mysids in a clade with Euphausiacea, Stomatopoda, and/or Syncarida. In contrast, phylogenomic studies (e.g., Schwentner et al. 2018; Höpel et al. 2022; Bernot et al. 2023; Yu et al. 2024; Dreyer et al. 2025) have consistently recovered a monophyletic Peracarida. However, most of these studies focused on broader pancrustacean relationships and had substantial gaps in peracarid taxon sampling; in particular, the above-mentioned relict lineages, except for Lophogastrida and/or Stygiomysida, were missing.

Central to this debate is the unique reproductive strategy of Peracarida that involves their ventral marsupium, where embryos develop directly into juvenile forms without free-living larval stages, although in two parasitic groups of isopods (Epicaridea and Gnathiidae) there are dispersive larval stages (e.g., Williams & Boyko 2012; Baeza 2015; Schädel et al. 2019; Williams et al. 2022). Peracarid direct development contrasts with that of other malacostracan crustaceans, such as decapods, which typically have complex dispersive larval phases (Martin et al. 2014). A striking exception to the ventral placement of the marsupium occurs in Thermosbaenacea, which brood their young in a dorsal carapace chamber (Olesen et al. 2015). Based on this character, it has been proposed that Thermosbaenacea are a sister group to the remaining Peracarida rather than part of Peracarida (i.e., Siewing, 1956, 1958, 1963; Richter & Scholz, 2001). However, most contemporary studies place Thermosbaenacea nested within Peracarida (e.g., Wagner 1994; Wills 1998; Poore 2005; Wilson 2009; Wirkner & Richter 2010). Notably, no phylogenomic study to date has included Thermosbaenacea, leaving this fundamental evolutionary question unresolved.

While most peracarid groups release juveniles that look like miniature adults (“manca” stages or similar), Mysidacea (Mysida, Stygiomysida, and Lophogastrida) produce an intermediate nauplioid that completes development within the pouch (pseudodirect development; San Vincente et al. 2014; Jirikowski et al. 2015). The monophyly of Mysidacea remains one of the most contentious questions within Peracarida. While morphological studies support its monophyly (Richter & Scholtz 2001; Poore 2005; Wirkner & Richter 2010; Grams et al. 2025), molecular analyses based on 18S or 28S rDNA reject this hypothesis, most notably by recovering Mysida outside of Peracarida and phylogenetically distinct from Lophogastrida and Stygiomysida (Jarman et al. 2000; Spears et al. 2005; Meland & Willasen 2007).

Over the years, various morphological studies have supported a sister group relationship of the two most diverse taxa, Amphipoda and Isopoda, based on shared characteristics like sessile eyes and a reduced carapace (e.g., Wagner 1994; Wills 1998; Poore 2005). However, this view has been challenged by both morphological and molecular studies that instead recover a clade uniting Isopoda with Cumacea and Tanaidacea - the Mancoida, named for their shared ‘manca’ larval stage (e.g., Siewing 1956; Richter & Scholtz 2001; Schwentner et al. 2018). Solving the phylogenetic positions of Thermosbaenacea, the relationships of Mysidacea, and testing the Mancoida hypotheses are a prerequisite to understanding the evolutionary transformations leading to broodcare within a marsupium and direct developing, non-dispersive larvae.

To resolve these crucial questions of peracarid evolution, we employed transcriptome and genomic data in a comprehensive phylogenomic approach designed to mitigate systematic error from heterogeneous rates of molecular evolution and compositional bias. We generated a densely sampled dataset of 156 malacostracans, with a core focus on 128 peracarids, including all orders except Bochusacea. This includes a comprehensive representation of Mysidacea, as well as the first phylogenomic data of the relict taxa Thermosbaenacea, Mictacea, Spelaeogriphacea, and Ingolfiellida. In addition, our expansive sampling within Amphipoda, Cumacea, Isopoda, Mysida, and Tanaidacea provides a robust higher-level phylogenetic framework for each of these species-rich taxa. These data were analyzed under multiple analytical frameworks and evolutionary models to ensure robust and congruent phylogenetic inference.

## Results

Our phylogenomic workflow incorporated genomic and transcriptomic data from 156 taxa, comprising 92 publicly available datasets (NCBI SRA; transcriptomes and genomes) and 64 newly sequenced transcriptomes, representing 128 peracarid and 28 other malacostracan taxa, representing a broad coverage of all non-peracaridan orders, for testing peracarid monophyly and the potential sister group of Peracarida. From these data, we constructed four complementary phylogenomic matrices: (1) a dataset based on Arthropoda BUSCO genes (673 loci; concatenated alignment length = 271,685 AA; 24% missing data), (2) a dataset with de novo orthogroup identification (ORTHO dataset: 511 loci; 189,938 AA; 29.7% missing data), and their respective signal-optimized subsets (3) BUSCO-50% (337 loci; 172,103 AA; 25.6% missing data) and (4) ORTHO-50% (255 loci; 95,481 AA; 29% missing data), created by retaining the top 50% most phylogenetically informative loci from each parent dataset. Phylogenetic reconstruction employed maximum likelihood (ML), Bayesian inference (BI), maximum parsimony (MP), and coalescent methods for a total of six approaches: ML in IQ-Tree2 with (i) ModelFinder used to find the best-fitting model for each gene, (ii) GHOST edge-unlinked mixture model, which accounts for heterotachous evolution, and (iii) PMSF site-specific frequency model, (iv) ASTRAL-III coalescence, (v) RAxML analysis (PROTOGAMMAAUTO model) of Dayhoff-recoded matrices, and (vi) MP in TNT. Notably, the TNT analysis was not conducted for the full BUSCO dataset due to time constraints and high computational demands. Bayesian inference (PhyloBayes MPI) was conducted on 50% datasets with a reduced taxon set (n = 62 taxa) with extended MCMC chains (>10,000 generations). We rooted all trees with Leptostraca, which is the sister group to all remaining Malacostraca (Richter & Scholtz 2001; Schwentner et al. 2018; Bernot et al. 2023; Dreyer et al. 2025).

All datasets and analytical methods congruently supported the monophyly of Peracarida and all peracarid orders with strong support (Fig.2). Within Peracarida, Mysidacea (Mysida + Lophogastrida + Stygiomysida) formed a monophyletic group with strong support; only some ASTRAL analyses had lower support for Mysidacea. Notably, the ASTRAL analysis of the ORTHO dataset failed to resolve deep peracarid relationships, resulting in a polytomy at the ordinal level. The majority of analyses recovered Mysidacea as the sister group to all other Peracarida; however, support for this placement varied. This was largely due to the unstable position of Thermosbaenacea, which was either recovered as the taxon sister to (1) Amphipoda + Ingolfiellida (the most commonly retrieved position), (2) all non-Mysidacea Peracarida, or (3) all other Peracarida (Fig.2). The latter was only recovered in the PhyloBayes analyses (with medium to strong). Amphipoda and Ingolfiellida were recovered as sister groups in all analyses, with strong to maximal support. Mancoida (Isopoda + Cumacea + Tanaidacea) likewise received strong support in all analyses. Within Mancoida, Isopoda and Cumacea were recovered as sister groups in most analyses with strong support, except for some ASTRAL and TNT analyses where Isopoda formed a clade with Tanaidacea, albeit with moderate support. Panmancoid (Mancoida + Mictacea + Spelaeogriphacea) formed a clade with high support in every analysis. However, internal relationships of this clade were inconsistent among analyses, with Mictacea recovered as the sister group to Mancoida or to Spelaeogriphacea (the latter mainly recovered in six-state recoding and TNT analyses).

**Figure 1.**
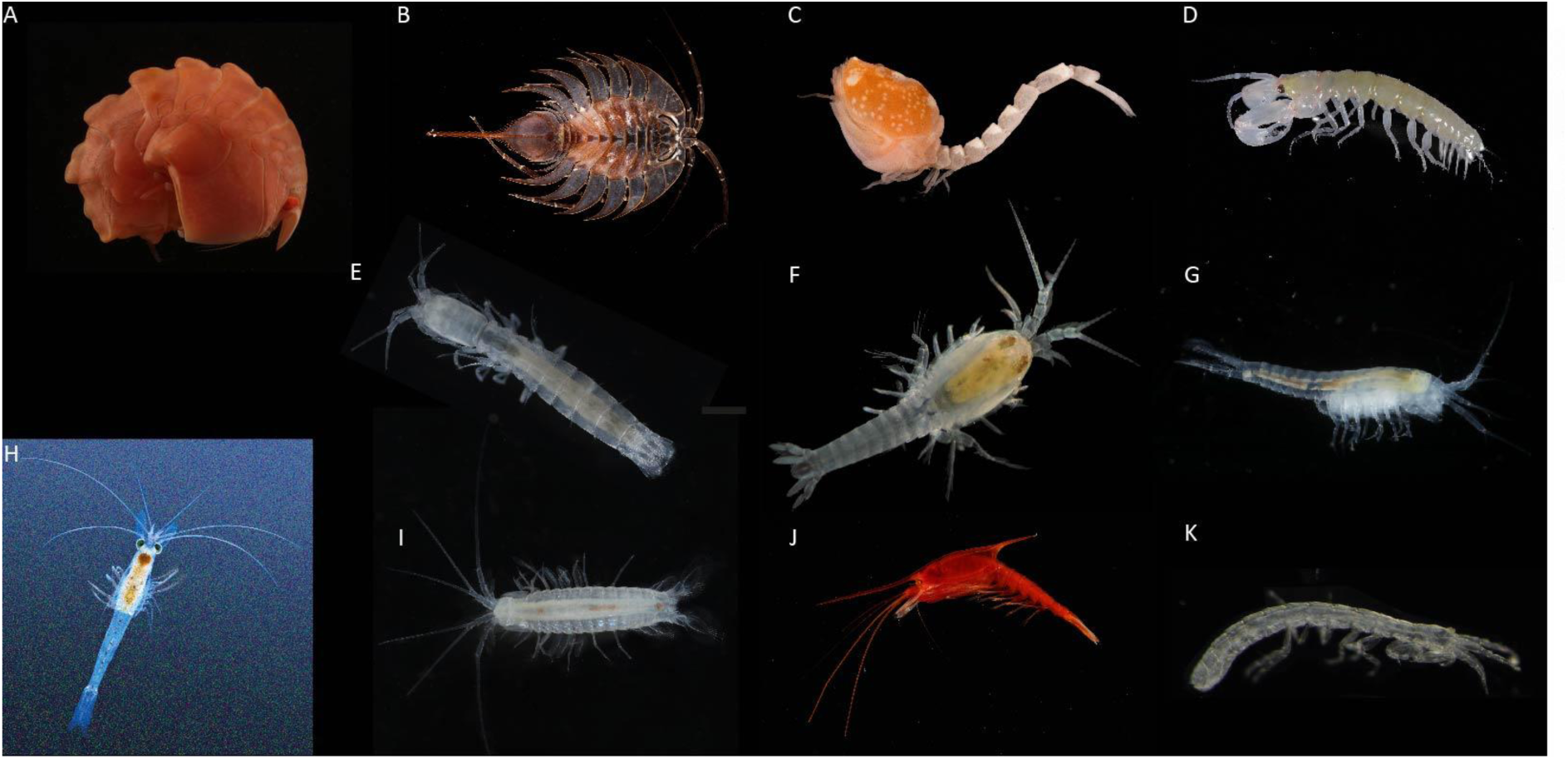
Morphological diversity of the studied orders within Peracarida. (A) *Epimeria inermis* (Amphipoda); (B) *Ceratoserolis meridionalis* (Isopoda); (C) *Cyclaspis gigas* (Cumacea); (D) Leptocheliidae (Tanaidacea); (E) *Stygiomysis clarkei* (Stygiomysida); (F) *Tulumella unidens* (Thermosbaenacea); (G) *Mictocaris halope* (Mictacea); (H) *Mysis relicta* (Mysida); (I) *Potiicoara brasiliensis* (Spelaeogriphacea); (J) *Gnathophausia zoea* (Lophogastrida); (K) *Ingolfiella longipes* (Ingolfiellida). Image credits: (A, B) Kevin Kocot, (C) Victoria VanderBrandt, (D) Kamila Głuchowska, (E, F, G, I, K) Jørgen Olesen, (H, J) Kenneth Meland

**Figure 2.**
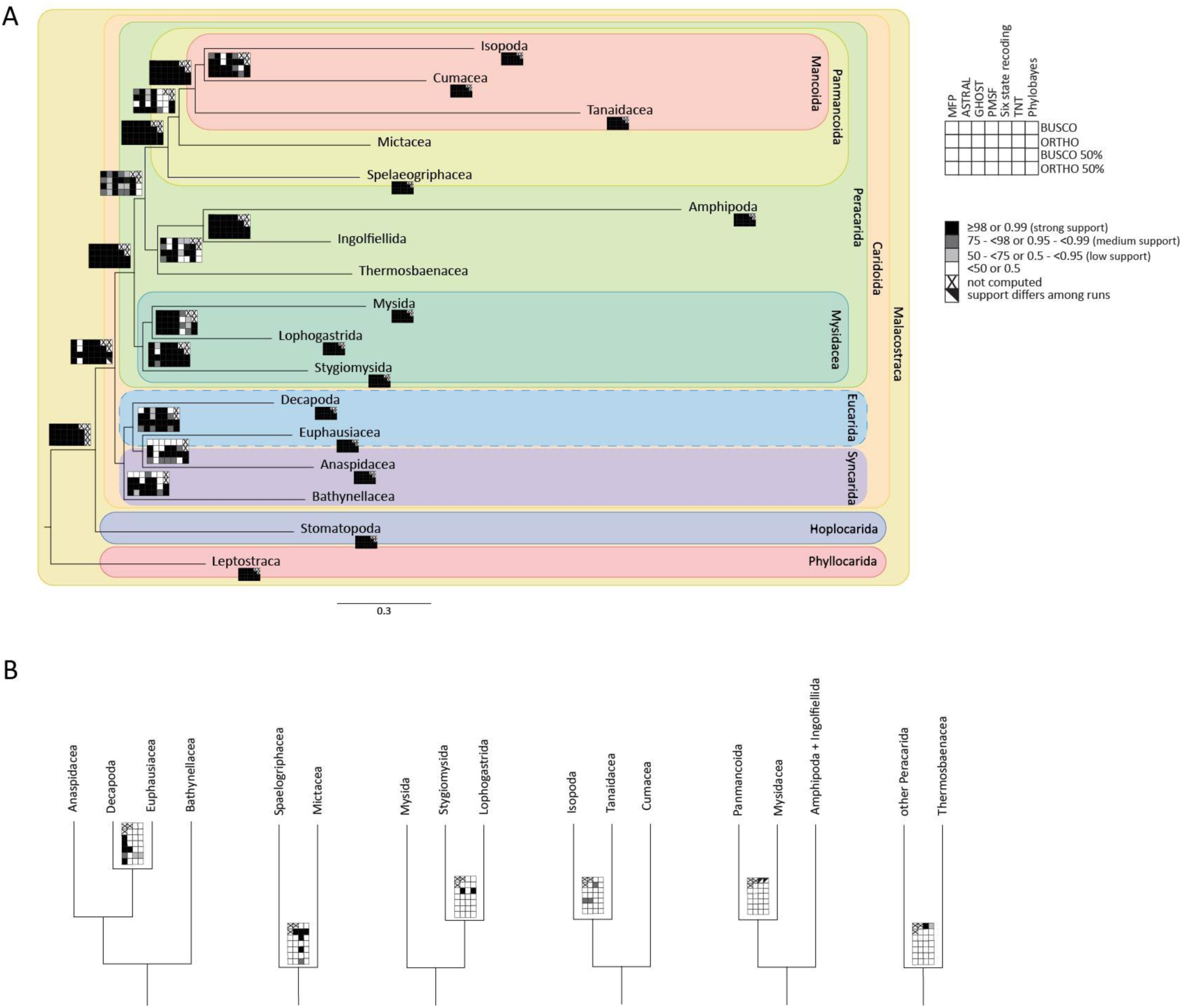
Summary of phylogenetic analyses. (A) Topology based on ORTHO-50% matrix comprising 255 decisive orthogroups and 95,481 amino acid positions analyzed with IQ-TREE MFP. Support values for all analyses are plotted on or below respective branches as specified in the legend at the top-right corner. Nodes are collapsed to show only the respective orders (matrix of support values are shown under each order name). Detailed relationships within orders are depicted in Supplementary Figures S2-S8 and all original trees are available at https://doi.org/10.57756/xanenh [this will become active after publication]; (B) Relevant alternative topologies.

Within Isopoda, our results robustly place Asellota as the sister group to all other isopods (Supplementary Fig.S5+S6). The subsequent divergence involves Gnathiidae (Cymothoida) and Epicaridea, which, form a clade in some analyses (Supplementary Fig.S5) but emerge as separate branches in others (Supplementary Fig.S6), consistently resulting in a paraphyletic Cymothoida. Oniscidea usually is the sister to all remaining Isopoda, followed by *Accalathura gigantissima* (Cymothoida); however, in some analyses, including all ORTHO-50% analyses, *A. gigantissima* diverges before Oniscidea. The relationships among the remaining sampled isopods are ambiguous. In the most commonly obtained topology, Limnoriidea branches off first, followed by the remaining Cymothoida (*Aegiochus* sp., Aegidae, *Cymothoa exigua*, *Aega* sp., *Bathynomus doederleinii*), and paraphyletic Sphaeromatidea (*Acutiserolis gerlachei*, *Gnorimosphaeroma oregonensis*, *Sphaeroma terebrans*) with regard to Valvifera (Supplementary Fig.S5). Notably, *Acutiserolis gerlachei* (Sphaeromatidea) consistently groups with Valvifera with high support in all analyses.

Within Cumacea, Diastylidae was consistently recovered as the sister taxon to all other Cumacea, with Nannastacidae as the sister taxon of Bodotriidae + Leuconidae with strong support (Supplementary Fig.S4). Notably, Bodotriidae was consistently resolved as paraphyletic with regard to Leuconidae, usually with strong support.

All tanaidacean superfamilies were recovered monophyletic with strong support Supplementary Fig.S8). Apseudomorpha was robustly placed as the sister group to the suborder Tanaidomorpha. Within Tanaidomorpha, the superfamily Neotanaoidea (represented by *Neotanais* cf. *kuroshio*) was consistently recovered as the sister taxon to Tanaidoidea. A notable inconsistency with the established systematics was found within Paratanaoidea, where *Parakanthophoreus* sp. (Akanthophoreidae) grouped with *Siphonolabrum* sp. and *Tanaella kommritzia* with strong support, and not with other Akanthophoreidae species (i.e., *Akanthophoreus* sp. and *Chauliopleona* cf. *sinusa*).

Within Amphipoda, some relationships were consistently recovered; however, the backbone within the order remained poorly resolved, and the position of several key taxa varied across analyses (Supplementary Fig.S2+S3). This may be largely due to the instability of Hyperiidea (represented by *Hyperia* sp.), which was recovered as either the most basal amphipod branch (Supplementary Fig.S3) or within a clade containing calliopioid, iphimedioid, and amphilochioid taxa (Supplementary Fig.S2). The subordinal classification was not supported. Senticaudata and Amphilochidea were found to be paraphyletic in all analyses. Talitroidea was monophyletic and consistently formed a clade with representatives of Hyloidea (*Hyalella azteca* and *Parhyale hawaiensis*), rendering the latter paraphyletic. The two Hadzioidea species formed a monophyletic group in the majority of analyses (Supplementary Fig.S3). A clade consisting of a monophyletic Calliopioidea, Iphimedioidea (*Sicafodia iceage*), and Amphilochoidea (*Neopleustes boecki*) was consistently recovered. The positions of lysianassoid taxa were highly variable. *Bathycallisoma schellenbergi* (Lysianassoidea) was often associated with the calliopioid-iphimedioid-amphilochoid clade (Supplementary Fig.S2) and was recovered with other lysianassoids in only a single analysis (Supplementary Fig.S3). While the infraorder Corophiida was monophyletic, the photoid *Jassa borowskyae* clustered with the Caprelloidea, rendering the latter paraphyletic. Gammaroidea formed a well-supported clade in all analyses (only *Eogammarus possjeticus* consistently grouped outside), with *Niphargus* sp. (Crangonyctoidea) as the sister group.

Within Mysidacea, Lophogastrida was recovered as the sister group to Mysida with strong support, except in the TNT analyses, where support was moderate to low. Analyses of the Dayhoff-recoded ORTHO and ORTHO-50% matrices, however, suggest a closer relationship between Lophogastrida and Stygiomysida, each with high support (only the ASTRAL ORTHO analysis resulted in a polytomy of Stygiomysida, Mysida + Lophogastrida, and the other Peracarida). Relationships within Mysida were stable and highly supported across all analyses (Supplementary Fig.S7). Petalophthalmidae (*Hansenomysis* sp.) was consistently recovered as the sister group to all other sampled Mysida (representing the family Mysidae).

Stomatopoda emerged as the sister group to Caridoida (Syncarida + Eucarida + Peracarida) in most analyses (Fig. 2), with high support (BS ≥ 98; PP ≥ 0.99), only some ASTRAL and in one chain of PhyloBayes ORTHO-50% topologies clustered Stomatopoda together with Syncarida and Eucarida.

Evolutionary rate comparisons revealed that Peracarida evolved significantly faster than their potential malacostracan sister gorup across all 25 phylogenetic analyses (p < 0.001 for all comparisons). A detailed examination of the ORTHO-50% matrix (analyses IQ-TREE, GHOST, PMSF) further revealed that this acceleration is a universal feature across all peracaridan orders. Pairwise t-tests showed that every peracaridan order (with n >3 taxa) also exhibited a significantly higher evolutionary rate than the non-peracaridan malacostracan sister group (p < 0.0001 for all; Supplementary Table S9). The IQ-TREE analysis of the ORTHO-50% matrix quantified these differences: the average branch length for Peracarida was 1.6 times longer than that the malacostracan sister group (Fig. 3). Among peracarids (n > 3), Amphipoda displays the highest evolutionary rate (1.84x), followed by Tanaidacea (1.72x). Mysida exhibits the lowest evolutionary rate of all peracarid orders (1.24x). The mean branch lengths for all groups are provided in Supplementary Table S10.

**Figure 3.**
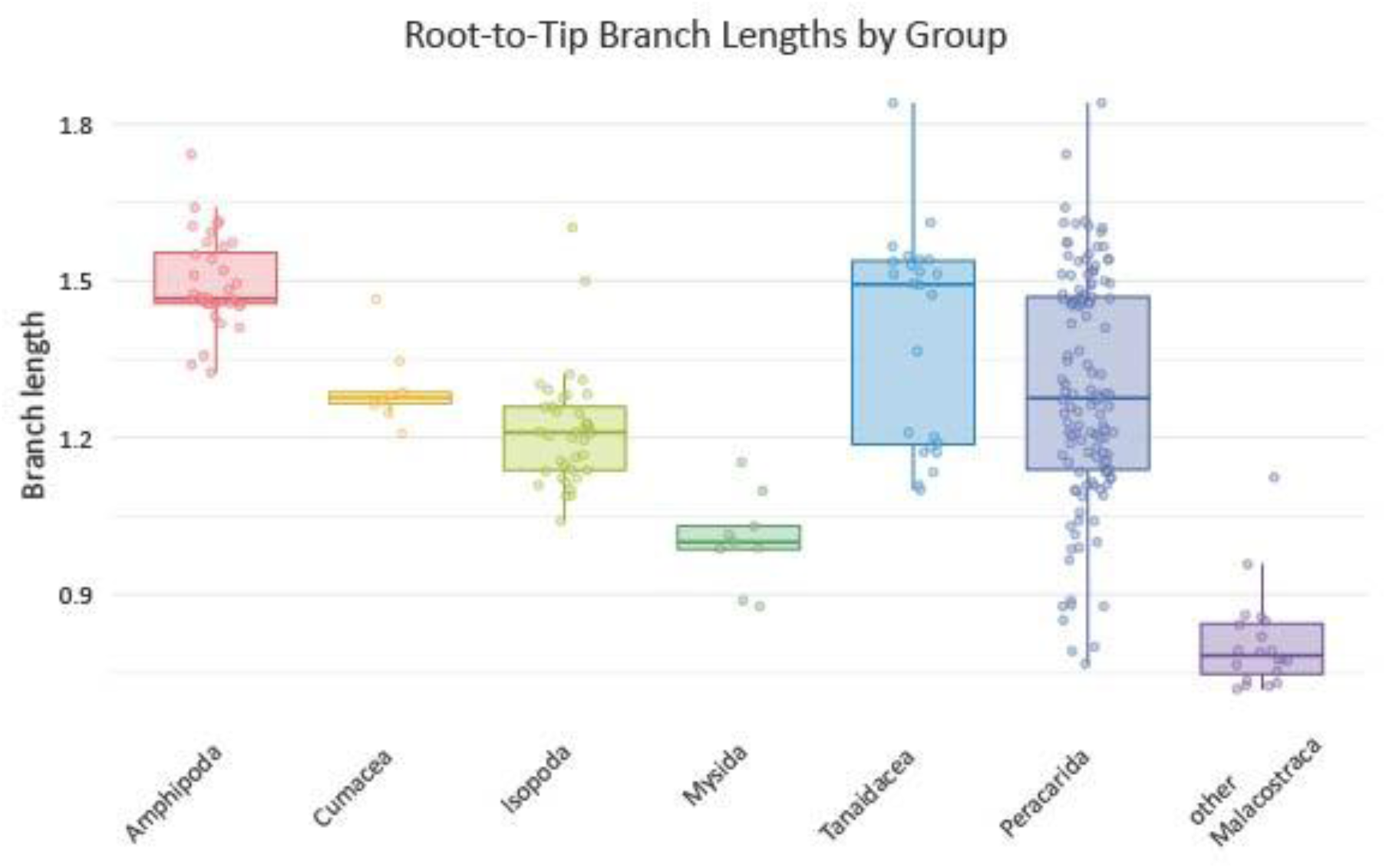
Branch length distribution of peracarid taxa and other Malacostraca. Boxplots show the distribution of root-to-tip branch lengths for five peracarid orders and for the combined groups “Peracarida” and “other Malacostraca”. Each box represents the interquartile range, with the horizontal line indicating the median and whiskers extending 1.5 times the interquartile range. Individual data points represent branch lengths calculated from individual gene trees of the ORTHO-50% matrix analyzed with IQ-TREE MFP and rooted with the three *Nebalia* species as the outgroup.

## Discussion

This study, covering 128 peracarid and 28 non-peracarid taxa and 255-673 loci per matrix, robustly recovered Peracarida as a monophyletic group, resolving a key controversy surrounding one of the most diverse and ecologically significant crustacean lineages. This result is consistent with previous morphological studies (e.g., Richter & Scholtz 2001; Poore 2005; Wirkner & Richter 2010; Grams et al. 2025) as well as phylogenomic analyses (e.g., Schwentner et al. 2018; Lozano-Fernandez et al. 2019; Höpel et al. 2022; Bernot et al. 2023; Dreyer et al. 2025). However, earlier phylogenomic analyses omitted several rare relict taxa, Thermosbaenacea, Mictacea, Ingolfiellida, and Spelaeogriphacea, which are crucial for a comprehensive understanding of peracarid phylogeny and were incorporated in our analyses. In contrast to earlier single-gene studies (e.g., Jarman et al., 2000; Spears et al. 2005; Meland & Willassen 2007) and some morphological studies (e.g., Watling 1991), that refuted peracarid monophyly, our results firmly place Mysida within monophyletic Peracarida and within monophyletic Mysidacea, rejecting a closer relationship of Mysida and Euphausiacea.

The evolution of brood care within a marsupium in Peracarida represents a major transition from the ancestral malacostracan life history, which typically includes a free-swimming larval stage that facilitates dispersal. Peracarids have secondarily lost this dispersive phase, with juveniles developing directly within a marsupium for nearly all groups (aside from epicaridean and gnathiid isopods). This is exemplified by the ‘manca’ stage of Mancoida, which highly resembles the adult. Our phylogeny suggests this transition potentially occurred via an intermediate step. Mysidacea hatch as a nauplioid larva that completes its development within the marsupium, a strategy termed ‘pseudodirect development’ (Jirikowski et al. 2015). The position of Mysidacea as the sister group to all remaining Peracarida implies that this may have been the ancestral peracarid condition from which the direct development evolved. Epicaridean and gnathiid isopods may provide an example of reacquisition (secondary gain) or perhaps evolutionary reversal to a free-swimming larval stage within the peracarids.

Resolving the position of Thermosbaenacea is fundamental for understanding the evolution of the peracarid marsupium, which is closely associated with extensive brood-care and (pseudo-)direct development. By including this pivotal taxon in a phylogenomic framework, our study reveals a critical conflict: our most complex Bayesian analyses place Thermosbaenacea as sister group to all other Peracarida (Siewing 1956,1963); all other analytical methods recover it nested within Peracarida. An ingroup peracarid position is supported not only by most of our analyses but also by numerous morphological studies (Wagner 1994; Wills 1998; Wilson 2009; Wirkner & Richter 2010, Grams et al. 2025), as well as earlier single-gene analyses (Spears et al. 2005; Meland & Willassen 2007) and is thus the current state of the art phylogenetic hypothesis. Notably, after our analyses were completed, the genome of *Tethysbaena scabra* was published (Pons et al. 2025). Future studies incorporating additional genomic data from Thermosbaenacea may help stabilize their position. However, the placement of Thermosbaenacea within Peracarida implies that their dorsal brood chamber is interpreted as a derived modification of ventral brooding, involving a marsupium formed by oostegites present in a common peracarid ancestor, or, alternatively, from a more simple ventral brood system, possibly involving egg pouches attached to the sixth thoracopods, as those seen in *Thermosbaena mirabilis* (Thermosbaenacea) where early embryos are carried prior to their relocation to the dorsal brood chamber (Zilch 1972; Olesen et al. 2015). This alternative, however, would be in conflict with the homology of oostegites.

Building on this evolutionary change in developmental mode, our phylogenetic results also reveal a corresponding pattern in molecular evolution: peracarid lineages consistently exhibit significantly longer branch lengths and thus higher rates of molecular evolution compared to other malacostracans. Notably, Mysida, which exhibit a pseudodirect development, potentially an intermediate evolutionary step, display the smallest branch lengths within Peracarida, reinforcing this correlation. Such long branches for Peracarida are not unique to our study, but can also be observed in other phylogenetic analyses (Spears 2005; Meland & Willasen 2007; Schwentner et al. 2018; Bernot et al. 2023; Yu et al. 2024; Dreyer et al. 2025). This accelerated rate of molecular evolution could be intrinsically linked to the evolutionary shift in their reproductive strategy as well as the associated changes in selection pressures and population genetic properties. The evolution of the marsupium and the loss of free-living larval stages transitioned most peracarids from an R-selected strategy toward a more strongly K-selected strategy, characterized by reduced offspring number and more limited dispersal capacity. This shift likely promotes population fragmentation and inbreeding, reducing the effective population size (N_e_). As outlined by Foltz (2003) and Bromham (2011), a smaller N_e_ enhances the effects of genetic drift, increasing the fixation rate of slightly deleterious mutations and thereby increasing the overall rate of molecular evolution.

Beyond these patterns of molecular evolution, our phylogenomic results also provide clarity on several deep phylogenetic relationships. Among these, we recover a notable congruence with major morphological hypotheses, recovering a well-supported clade uniting Mictacea, Spelaeogriphacea, and Mancoida. The robustness of this clade is further reinforced by comprehensive morphological data (Grams et al. 2025; Richter & Scholz 2001; Wirkner & Richter 2010), which placed Bochusacea within this clade as well (Wilson 2009; Grams et al. 2025). Some earlier studies had even included Spelaeogriphacea in Mancoida (Watling 1981). In fact, manca larva, charakteristic of mancoid taxa, have been reported for Spelaeogriphacea (Poore & Humphreys 2003; Pires-Vanin 2012), Mictacea (Bowman & Iliffe, 1985) and one species of Bochusacea (Gutu et al. 1998). For consistency, we suggest retaining the taxon Mancoida exclusively for the taxa tracing back to the last common ancestor of Cumacea, Isopoda, and Tanaidacea (which might also include Bochusacea; Wilson 2009; Grams et al. 2025). Within Mancoida, Tanaidacea forms the most basal branch, and Isopoda and Cumacea are strongly supported as sister groups (see also Meland & Willassen 2007; Wirkner & Richter 2010; Bernot et al. 2023). We herein propose the new taxon name **Panmancoida new taxon** (derived from “pan” [Greek: “all” or “entire”] and Mancoida, which refers to the manca larva) for the extended taxon including Mancoida, Mictacea, Speleogriphacea, and Bochusacea. The monophyly of Panmancoida is also supported by morphological data. A synapomorphy of Panmancoida is an early embryo with dorsally flexed caudal papillae (Richter & Scholtz 2001; Olesen et al. 2015).

According to the classification of Lowry & Myers (2017), Ingolfiellidea was placed outside of Amphipoda and raised to order status. Our analysis placed Amphipoda as the sister group to Ingolfiellida. Hence, classifying them as separate orders or uniting them within a broader concept of Amphipoda are both justified interpretations. Some morphological studies supported a sister-group relationship between the species-rich taxa Isopoda and Amphipoda (at that time including Ingolfiellida), based on putative synapomorphies such as sessile eyes, the reduction of the carapace, and the reduction of exopods on the thoracopods (e.g., Wagner 1994; Wills 1998; Poore 2005; Grams et al. 2025). However, our recovery of Mancoida is in line with other morphological and molecular studies (e.g., Richter & Scholtz 2001; Wilson 2009; Wirkner & Richter 2010; Spears et al. 2005; Meland & Willassen 2007; Schwentner et al. 2018; Höpel et al. 2022; Bernot et al. 2023) rejecting a closer relationship of Amphipoda and Isopoda.

Our phylogeny of Isopoda consistently recovers monophyly of the major suborders Valvifera, Oniscidea, and Asellota, which is well supported by molecular and morphological work (Wägele 1989; Wetzer et al. 1997; Thomas Thorpe 2024). Furthermore, our findings strongly support the ‘CLVS’ clade (Cymothooidea, Limnoriidea, Valvifera, Sphaeromatidea) proposed by Thomas Thorpe (2024). Within this clade, our results suggest a sister-group relationship between Valvifera and Seroloidea (Sphaeromatidea), rendering Sphaeromatidea paraphyletic (see also Wetzer et al. 2013; Lins et al. 2017; Thomas Thorpe 2024). Notably, our analysis places Anthuroidea (Cymothoida) as the sister group to the entire CLVS clade, a finding that would render Cymothoida paraphyletic and suggests Anthuroidea may represent a distinct lineage. While current taxonomy places Gnathiidae within Cymothoida, our results suggest a closer relationship between Gnathiidae and Epicaridea (see also Brusca & Wilson 1991). The entire CLVS clade (excluding Gnathiidae) forms the sister group to Oniscidea, with Epicaridea, Gnathiidae, and Asellota occupying subsequent basal positions. Of course, one should keep in mind that some important isopod taxa like Phreatoicidea could not be included. Our findings, congruent with Thomas Thorpe (2024), demonstrate that parasitism evolved independently in Cymothoida and in the Epicaridea-Gnathiidae clade. The sister-group relationship between Epicaridea and Gnathiidae implies a hematophagous common ancestor, contradicting the hypothesis of an epicaridean origin from fish parasites (Dreyer & Wägele 2001). While these results clarify major evolutionary transitions, a broader sampling of parasitic taxa (e.g., Cirolanidae, Corallanidae) remains essential for a comprehensive understanding.

We present a well-supported phylogeny for Tanaidacea that corroborates the superfamilial relationships with current taxonomy: Apseudoidea represents the most basal branch, followed by the successive divergences of Paratanaoidea and a clade comprising the sister groups Tanaidoidea and Neotanaoidea. This topology is fully congruent with earlier findings of Drumm (2010), based on three single markers (COI, H3, and 28S), and is further corroborated by the transcriptome-based phylogeny of Kakui et al. (2021), from which we sourced the majority of our tanaidacean sequences.

Our analysis largely confirms the monophyly of major cumacean families (see also Gerken et al. 2022), though we recover Bodotriidae as paraphyletic. We strongly support the position of Diastylidae as the sister group to all other cumaceans included (Haye et al. 2004; Luque & Gerken 2019; Gerken et al. 2022). However, our results suggest a key revision of the internal relationships of the latter clade: we find robust support for a closer relationship between Bodotriidae and Nannastacidae, with Leuconidae representing their sister group. This stands in contrast to the topology of Gerken et al. (2022), which suggested Leuconidae and Nannastacidae as sister taxa. Our current limited taxon sampling highlights a need for future comprehensive phylogenies with denser sampling, including the missing cumacean families Ceratocumatidae, Gynodiastylidae, Lampropidae, and Pseudocumatidae, to fully resolve cumacean evolutionary relationships.

Our results highlight the unresolved backbone of amphipod phylogeny, contrasting with current taxonomic frameworks (Lowry & Myers 2017). Deep relationships, including the placement of Hyperiidea and the paraphyletic clades of Senticaudata and Amphilochidea, remained unstable (see also Copilaş-Ciocianu et al. 2020; Bernot et al. 2023; Dreyer et al. 2025). The paraphyly of Hyaloidea and Gammaroidea, as well as the variable positions of lysianassoid taxa, imply that diagnostic morphological characters may not reliably reflect evolutionary history in these groups. Overall, our findings emphasize the need for an expanded taxonomic sampling to stabilize deep nodes of the Amphipoda.

We robustly recover Mysida as sister group to Lophogastrida and Stygiomysida, and therefore a monophyletic Mysidacea (see also Höpel et al. 2022; Bernot et al. 2023; Grams et al. 2025).

Mysida comprises two families, historically differentiated by the presence (Mysidae) or absence (Petalophthalmidae) of a statocyst in the uropods (Meland et al. 2015). Our phylogeny robustly supports this morphological classification: *Hansenomysis* sp. (Petalophthalmidae) was recovered as the sister group to all other sampled mysids, representing 5 of 10 subfamilies in the family Mysidae (Meland & Willassen 2007; Meland et al. 2015; Kou et al. 2025).

The phylogenetic placement of Stomatopoda has been a persistent challenge in malacostracan systematics. Our analysis provides robust support for positioning Stomatopoda as the sister group to Caridoida (all other Eumalacostraca). This placement is consistent with a subset of recent phylogenomic analyses (Schwentner et al. 2018; Bernot et al. 2023) as well as previous morphological studies (e.g., Siewing 1956; Richter & Scholtz 2001; Wirkner & Richter 2010) and challenges the alternatively proposed closer relationship between Stomatopoda and Eucarida and/or Syncarida (e.g., Schwentner et al. 2018; Höpel et al. 2022; Bernot et al. 2023). Relationships among other major lineages remain contentious. The monophyly of Eucarida (Decapoda + Euphausiacea) is notably unresolved, as Euphausiacea forms a clade either with Decapoda or with Anaspidacea. Syncarida (Bathynellacea + Anaspidacea) is consistently recovered as polyphyletic, similar to the findings of Bernot et al. (2023), though the unstable placement of *Allobathynella bangokensis* (Bathynellacea) underscores lingering uncertainty in syncarid interrelationships.

This phylogenomic study provides a significantly resolved phylogenetic framework for Peracarida, one of the most species-rich taxa within crustaceans, reconciling long-standing conflicts between morphological and molecular data. Our robust support for a monophyletic Peracarida, with Stomatopoda as the sister group to all other Eumalacostraca (Malacostraca except Leptostraca), offers a stable foundation for understanding the evolution of this ecologically crucial group. We demonstrate that key peracarid innovations - the marsupium and direct development - drove a shift in reproductive strategy, reducing dispersal and effective population size. This shift may provide a mechanistic explanation for the accelerated molecular evolutionary rates observed across peracarid lineages, linking a profound change in developmental mode to its long-term genomic consequences. While deep relationships within some taxa like Amphipoda remain challenging, our results provide a backbone across Isopoda, Tanaidacea, Cumacea, and Mysidacea. This work underscores the necessity of combining dense taxon sampling with phylogenomic data to test morphological hypotheses and reconstruct the evolutionary history of complex radiations. Nonetheless, this study also charts a course for future research. Critical next steps include incorporating more Thermosbaenacea and representatives of Bochusacea to further test the relationships of Peracarida, especially concerning the contentious placement of Thermosbaenacea.

## Materials and Methods

### Sample collection and sequencing

Taxon sampling comprised 156 taxa represented by annotated genomes or transcriptomes. Of these, 43 publicly available transcriptomes downloaded from the NCBI Sequence Read Archive (SRA) as raw reads, 43 assembled transcriptomes were downloaded from NCBI TSA, and 6 genome annotations were downloaded from NCBI Genome (Supplementary Table S1). Publicly available data were supplemented with 64 transcriptomes that were newly sequenced for this study. Outgroup taxon sampling was performed to span all extant non-peracarid malacostracan lineages, including species spanning the phylogenetic diversity of each clade. Leptostraca (*Nebalia* spp.) was used to root the trees.

For libraries generated by the Kocot lab (Supplementary Table S1), RNA was extracted from RNAlater or cryo-preserved specimens using the Omega Bio-Tek RNA MicroElute Kit. RNA concentration was measured using a Thermo Fisher Qubit 3.0 fluorometer with the RNA High Sensitivity kit, purity was assessed by measuring the 260/280 nm absorbance ratio using a Thermo Fisher Nanodrop Lite, and integrity was evaluated using a 1% SB agarose gel. Complementary DNA (cDNA) synthesis was performed using 1 ng of total RNA using the Clontech SMART-Seq HT kit. An Illumina sequencing library was then prepared using either the Takara SMART-Seq HT Plus kit with 2-10 ng of cDNA or the Nextera XT kit with 0.125 ng of cDNA. Complementary DNA and Sequencing library molecular weight distribution and concentration were assessed using an Agilent Fragment Analyzer with the NGS Fragment Kit (1-6000 bp). Resulting libraries were pooled and sent to Azenta (Plainsfield, NJ, USA) for sequencing on an Illumina NovaSeq S4 flowcell with 2 X 150 bp PE reads. For libraries generated by the Schwentner lab, whole specimens were preserved in RNAlater, stored at −80°C, and submitted to Macrogen (Amsterdam, Netherlands) for RNA extraction, cDNA synthesis, and sequencing library preparation using the TruSeq Stranded mRNA Library Prep kit. In some cases, the digestive tract was removed before preservation.

### Transcriptome Assembly and Phylogenomic Dataset Construction

Raw reads were assembled into transcripts using *Trinity v2.132 or v2.10.0* (Grabherr et al., 2011). Assembly completeness was assessed by using *BUSCO v5.2.2* (Manni et al. 2021) against the *arthropoda_odb10* database; assemblies with low *BUSCO* scores were excluded unless critical for taxonomic representation.

Coding regions were predicted using *TransDecoder v5.7.1* (Haas, BJ. https://github.com/TransDecoder/TransDecoder), initially retaining all open reading frames (ORFs) ≥ 100 amino acids (AA). To maximize sensitivity for functionally significant ORFs, we validated these predictions through: (1) *DIAMOND v2.1.9* (Buchfink et al. 2021) similarity searches against the *UniProtKB/Swis-Prot* database, and (2) domain detection using *HMMER v3.4* (Zhang & Wood 2003). The resulting BLAST and Pfam outputs were integrated through TransDecoder.Predict, with applied filters to retain only the single best coding region prediction per transcript meeting a minimum length threshold of 500 AA. For the ORTHO datasets, orthologous groups were inferred using *OrthoFinder v2.5.5* (Emms & Kelly 2019). To mitigate biases from gene duplication or misassembly, orthogroups containing >10x transcripts per taxon were excluded. Fragmentary sequences <100 AA were discarded. For the BUSCO datasets, single-copy orthologs identified by BUSCO were extracted using a custom Python script (BUSCO_retrieve_genes.py) to create the first phylogenomic dataset (= BUSCO).

Retained sequences from both datasets were aligned with *MAFFT v7.487* (Katoh & Standley 2013). The ORTHO dataset alignments were further cleaned with *HmmCleaner* (Di Franco et al. 2019) to remove potentially mistranslated sequence regions. Alignments sampled for <50% of taxa after this cleaning were discarded to minimize overall missing data. Gene trees were reconstructed for each orthogroup using *FastTree v2.2* (Price et al. 2009, 2010) with the -gamma model. Trees were processed with *PhyloPyPruner* (https://gitlab.com/fethalen/phylopypruner) to remove paralogous sequences, retaining only one sequence per taxon per orthogroup, while simultaneously applying a 75% minimum taxon occupancy threshold.

The BUSCO and ORTHO alignments were trimmed using *ClipKIT v2.3.0* (Steenwyk et al. 2020) to remove sites where gaps occurred in ≤50% of sequences. Trimmed alignments were filtered again using *PhlyoPyPruner* (75% occupancy). We retained 673 loci from the BUSCO dataset and 511 from the ORTHO dataset for downstream analyses. From these datasets, two additional datasets were created by filtering for the most informative loci. Using *genesortR* (Mongiardino Koch 2021), an R-based tool that quantifies phylogenetic utility through metrics such as parsimony-informative sites and branch support, all loci were ranked by their evolutionary informativeness. For each dataset, we retained only the top 50% highest-ranking loci, resulting in a BUSCO-50% dataset (337 loci) and an ORTHO-50% dataset (255 loci).

### Phylogenetic Analyses

We subjected all four datasets to six different phylogenetic approaches. We performed three maximum likelihood analyses in *IQ-TREE 2 v1.5.0* (Minh et al. 2020): (1) standard inference using the ModelFinder-embedded MFP algorithm, (2) the GHOST framework (Crotty et al. 2020), which partitions sites into multiple rate categories, and (3) the site-heterogeneous PMSF approximation (Wang et al. 2018), using posterior mean site frequencies derived from a profile mixture model. Branch support for all three analyses was assessed through 1000 ultrafast bootstrap replicates (UFBoot). We further implemented a coalescence-based approach using ASTRAL-III v5.7.7 (Zhang et al. 2018) to account for incomplete lineage sorting. Input gene trees were generated in *IQ-TREE 2* under the MFP model with 1000 UFBoot replicates to assess branch support. To mitigate potential artifacts from compositional heterogeneity and substitution saturation, we also implemented Dayhoff 6-state recoding, which clusters amino acids into six functionally similar groups based on substitution probabilities from the Dayhoff/Pam250 matrix (Dayhoff et al. 1978). The recoded alignments were then analyzed in *RAxML-NG v8.2.12* (Stamatakis 2014) using the MULTIGAMMAI model with 1000 UFBoot replicates to assess branch support. Finally, we conducted a maximum parsimony analysis using *wTNT v.1.6 2025-06-22* (Goloboff & Morales 2023) using the “PhylogenomicSearch.run” script (Torres et al. 2022) with the command “`run PhylogenomicSearch.run input DATASET_NAME.tnt search nt level 4 hits 2 output newick;`” and calculating bootstrap support with the “PhylogenomicSupport.run” script (Torres et al. 2022) with the command “`run PhylogenomicSupport.run input DATASET_NAME.tnt reftree REFERENCETREE_NAME.tre sites boot replic 1000 search nt level 1 output newick;`”. Due to time constraints and high computational demand, we were not able to conduct the TNT analysis for the full BUSCO dataset.

Representing the most complex model used in our study, Bayesian inference was implemented using *PhyloBayes MPI v1.9* (Lartillot et al. 2013) under the CAT-GTR model. Due to the method’s computational demands, we restricted this analysis to the 50% datasets (BUSCO-50% and ORTHO-50%) with a strategically reduced taxon set (n = 62), preserving all major clades. For the ORTHO 50% dataset, three independent chains were run for 10,069 generations each, with sampling every 10th generation after discarding 2,500 generations as burn-in. The BUSCO 50% dataset was analyzed with two chains of 10,202 generations under identical sampling and burn-in parameters. We discarded the first 2500 trees from each chain as burn-in and calculated a 50% majority rule consensus tree from the remaining trees from each chain.

### Evolutionary Rate Calculations

To quantify evolutionary rates across Malacostraca, we calculated root-to-tip branch lengths for each of the 25 phylogenetic analyses using custom Python and Bash scripts. All trees were rooted with Leptostraca as the outgroup to standardize distance measurements. We excluded *Hyperia* sp. due to its exceptionally long branches, which could disproportionately influence rate comparisons. We first used a t-test on each phylogenetic analysis to determine whether Peracarida, as a whole, exhibited significantly different evolutionary rates compared to their potential malacostracan sister group (comprising all other malacostracan lineages except Leptostraca and Stomatopoda). For a detailed investigation of the ORTHO-50% matrix (IQ-TREE), we calculated mean branch lengths for Peracarida, each peracaridan order, and the non-peracaridan sister group. Subsequently, on the ORTHO-50% matrix trees (IQ-TREE, GHOST, PMSF), we performed pairwise t-tests comparing each peracaridan order (with n > 3 taxa) against this non-peracaridan malacostracan group.

## Supporting information

Supplementry Figrue S2

Supplementry Figrue S3

Supplementry Figrue S4

Supplementry Figrue S5

Supplementry Figrue S6

Supplementry Figrue S7

Supplementry Figrue S8

## Author Contributions

Conceptualization, A-C.B., M.S., S.R., R.W., K.K., and S.G.; formal analyses, A-C.B., K.K., and M.G.; resources, A-C.B., M.G., H.B., S.B., L.M.C., B.C., S.C., W.J.F, S.G., C.G.H., A-N.L., S.M., K.M., J.O., P.A.P., S.R., R.W., J.W., K.M.K., and M.S.; visualization, A-C.B.; writing - original draft, A-C.B.; writing - review and editing, A-C.B., M.G., H.B., S.B., L.M.C., B.C., S.C., S.G., C.G.H., A-N.L., S.M., K.M., L.K., J.O., P.A.P., S.R., R.W., J.W., K.M.K., and M.S.

## Acknowledgments

We thank Martin Kapun for assisting during bioinformatic analyses. We further extend our gratitude to Jessica Thomas Thorpe, Lauren Hughes, and Magdalena Błażewicz for their valuable contributions in discussing our results, as well as George (Buz) Wilson for his detailed identifications of the isopod material included. Photographs used were kindly provided by Victoria VanderBrandt and Kamila Głuchowska. Karen Jeskulke, Carolin Uhlir, and Nicole Gatzemeier are thanked for curating all samples via the onboard databases during several expeditions, as well as supporting the sample transfer.

## Data Availability

Sample information and sequence data for de novo transcriptomes are available in NCBI [Bioproject will become active after publication]. Additional materials, including phylogenetic alignments, tree files, branch length metrics, and custom analysis scripts (Bash/Python), have been deposited in our data repository (https://doi.org/10.57756/xanenh) [will become active after publication].

## Supplementary Material

Supplementary Table S1. Overview of taxonomic sampling and data sources.

The table lists all taxa included in the analysis, their corresponding NCBI IDs, the source of the genomic or transcriptomic data, as well as BUSCO metrics of the assembled data.

Supplementary Figure S2. Detailed phylogenetic relationships within Amphipoda.

Support values for all analyses are plotted on respective branches as specified in the legend at the top-right corner. Topology is based on IQ-TREE MFP analysis of ORTHO-50% matrix. All original trees are available at https://doi.org/10.57756/xanenh [this will become active after publication].

Supplementary Figure S3. Detailed phylogenetic relationships within Amphipoda.

Support values for all analyses are plotted on respective branches as specified in the legend at the top-right corner. Topology is based on ASTRAL analysis of ORTHO-50% matrix. All original trees are available at https://doi.org/10.57756/xanenh [this will become active after publication].

Supplementary Figure S4. Detailed phylogenetic relationships within Cumacea.

Support values for all analyses are plotted on respective branches as specified in the legend at the top-right corner. Topology is based on IQ-TREE MFP analysis of ORTHO-50% matrix. All original trees are available at https://doi.org/10.57756/xanenh [this will become active after publication].

Supplementary Figure S5. Detailed phylogenetic relationships within Isopoda.

Support values for all analyses are plotted on respective branches as specified in the legend at the top-right corner. Topology is based on IQ-TREE MFP analysis of ORTHO-50% matrix. All original trees are available at https://doi.org/10.57756/xanenh [this will become active after publication].

Supplementary Figure S6. Detailed phylogenetic relationships within Isopoda.

Support values for all analyses are plotted on respective branches as specified in the legend at the top-right corner. Topology is based on PMSF analysis of BUSCO matrix. All original trees are available at https://doi.org/10.57756/xanenh [this will become active after publication].

Supplementary Figure S7. Detailed phylogenetic relationships within Mysidacea.

Support values for all analyses are plotted on respective branches as specified in the legend at the top-right corner. Topology is based on IQ-TREE MFP analysis of ORTHO-50% matrix. All original trees are available at https://doi.org/10.57756/xanenh [this will become active after publication].

Supplementary Figure S8. Detailed phylogenetic relationships within Tanaidacea.

Support values for all analyses are plotted on respective branches as specified in the legend at the top-right corner. Topology is based on IQ-TREE MFP analysis of ORTHO-50% matrix. All original trees are available at https://doi.org/10.57756/xanenh [this will become active after publication].

Supplementary Table S9. Results of paired t-tests comparing branch lengths among peracarid orders and other malacostracans.

Paired t-tests were conducted for thre phylogenomic analyses (IQ-TREE MFP, PMSF, GHOST) based on the ORTHO-50% matrix. The table lists t-vales, degrees of freedom and p-values for each comparison.

Supplementary Table S10. Mean branch lengths of IQ-TREE MPF analysis based on ORTHO-50% matrix.

Mean branch lengths are calculated for every peracarid order (n>3, excluding *Hyperia* sp.) and the remaining malacostracans (except Leptostraca and Stomatopoda).

