## Supplementary figures and images for "Phylogenomics supports monophyly of marsupial crustaceans: a journey to direct development"

### Supplementry Figrue S2

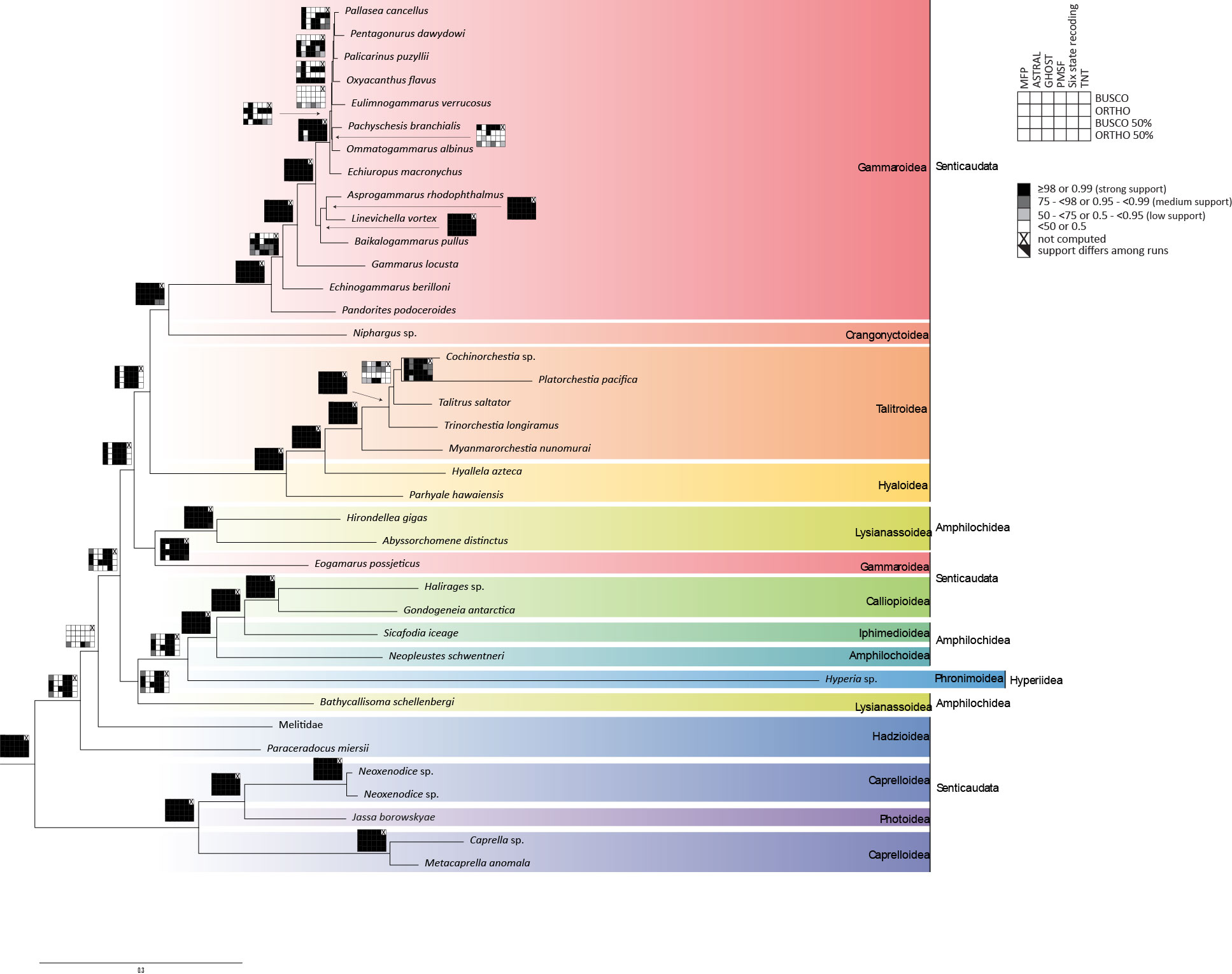

### Supplementry Figrue S3

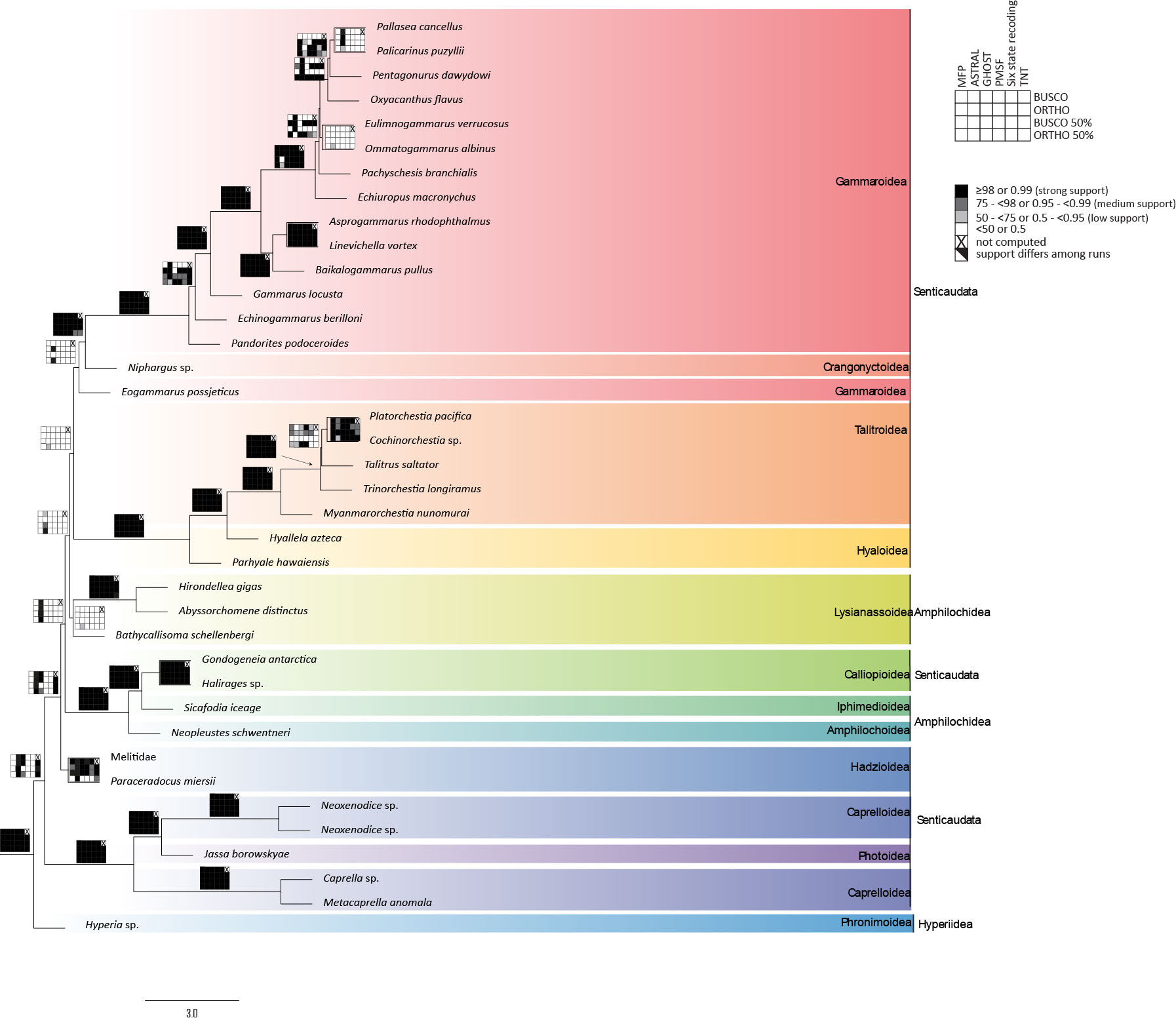

### Supplementry Figrue S4

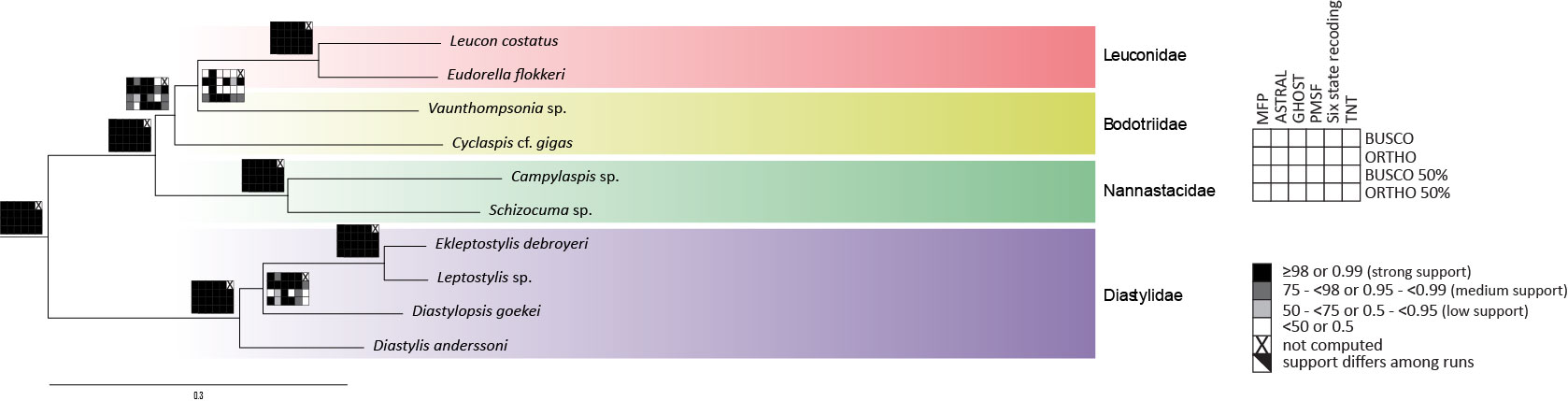

### Supplementry Figrue S5

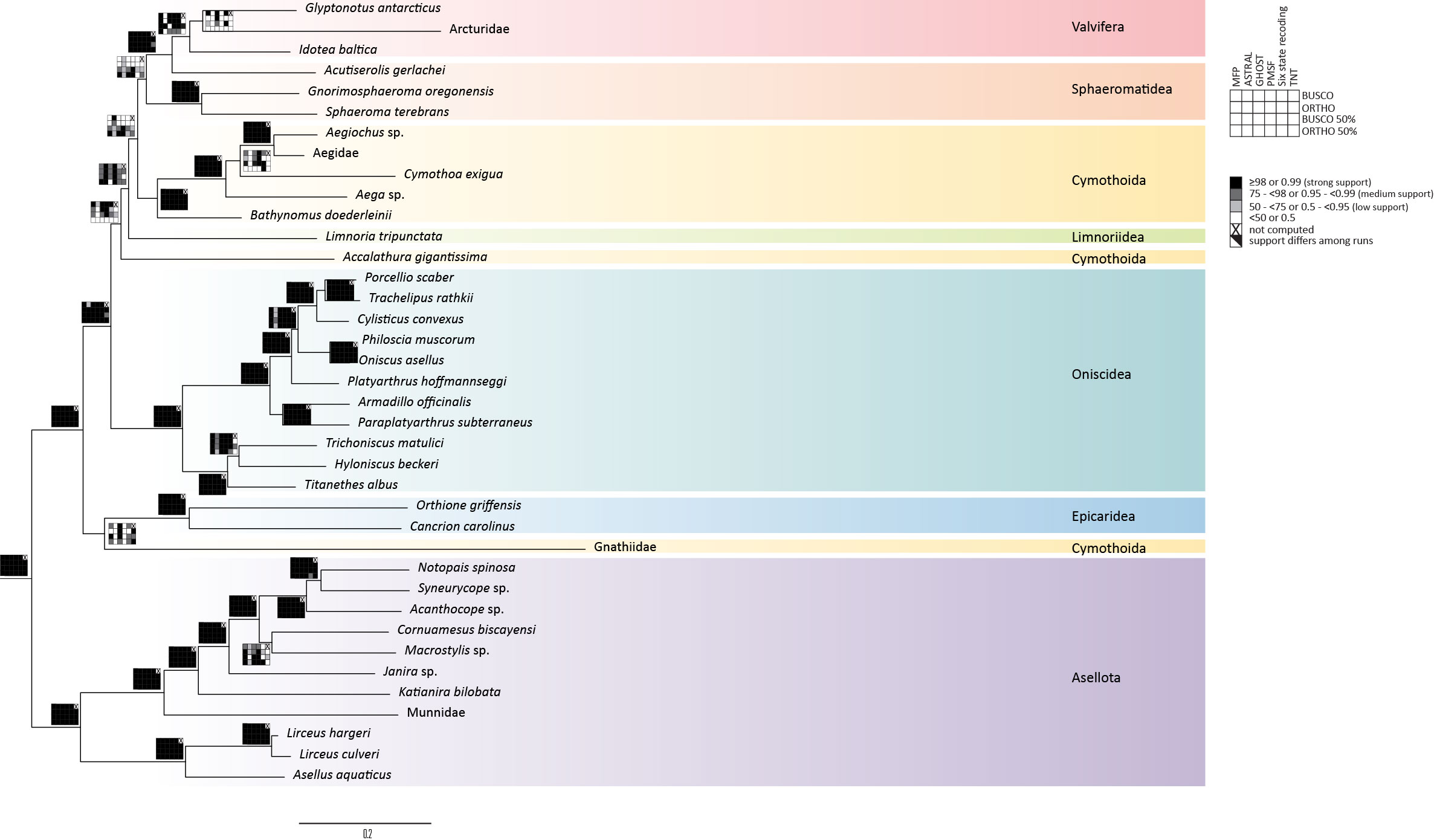

### Supplementry Figrue S6

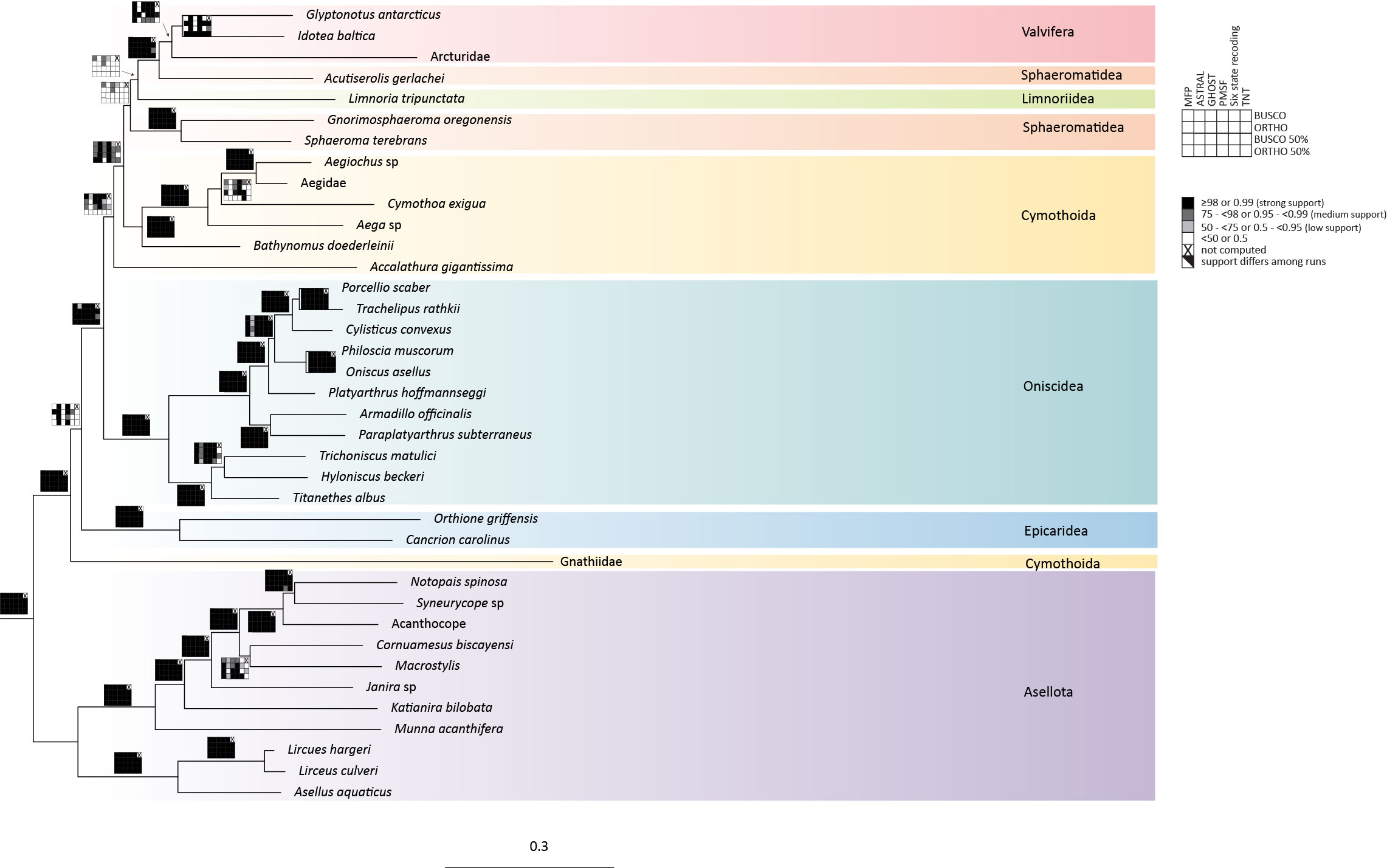

### Supplementry Figrue S7

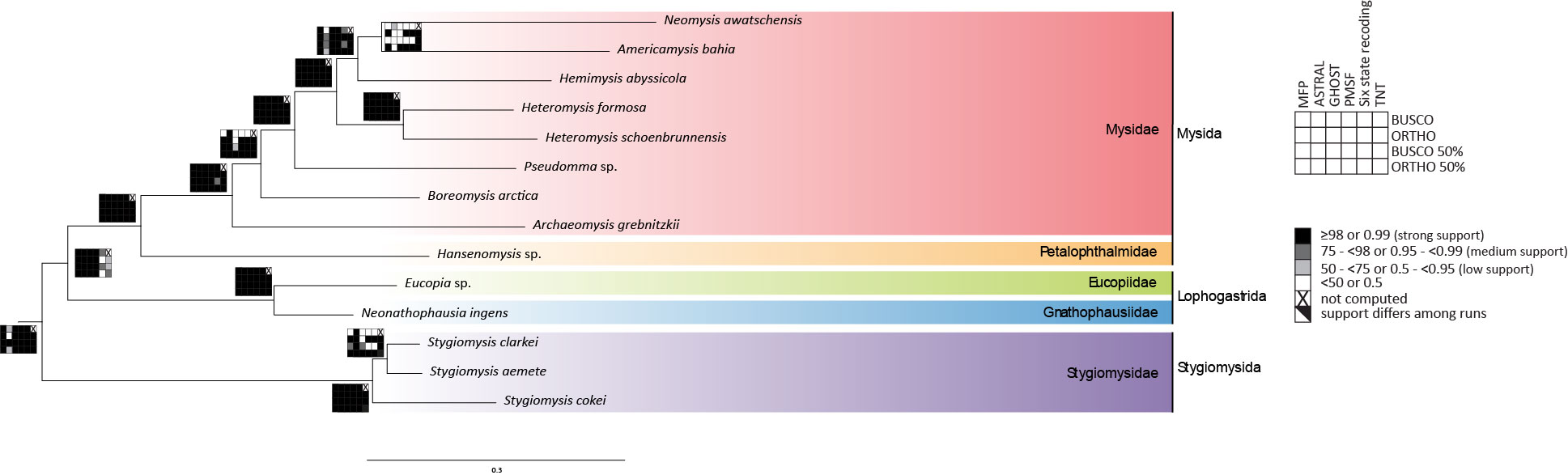

### Supplementry Figrue S8

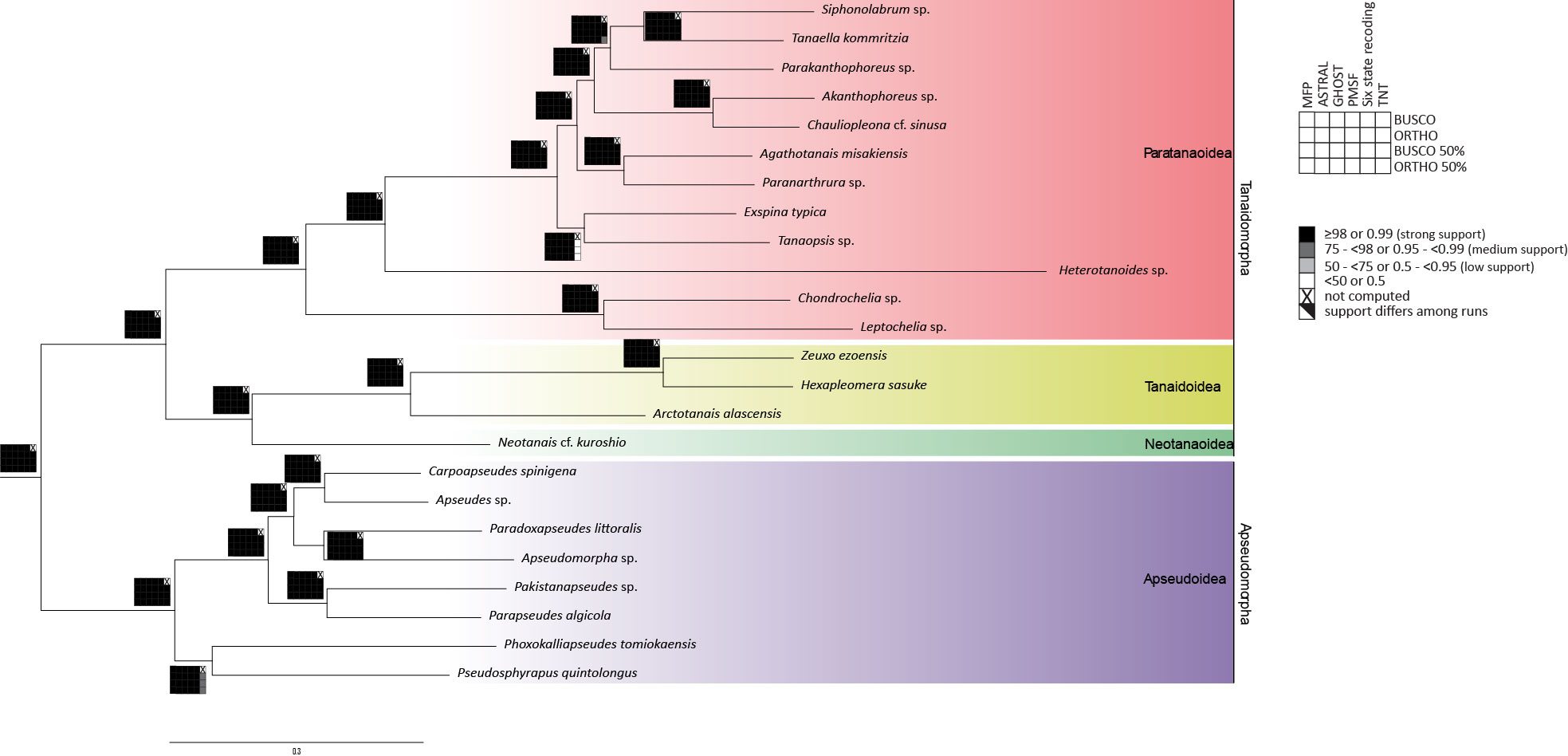
